# EXtrACtor – A tool for multiple queries and data extractions from the EXAC and gnomAD database

**DOI:** 10.1101/483909

**Authors:** Flip Mulder, Bobby P.C. Koeleman

## Abstract

**Introduction:** EXtrACtor provides an easy and flexible way to perform multiple queries of the ExAC and gnomAD databases (Lek M, al. e, 2016) and process the results. The ExAC browser is an often-used tool to gather information from a large number of exomes while the gnomAD database expands on this by increasing the number of exomes but also adding genomes. However the websites only allow the users to query a single gene, transcript or location at a time. On the other hand downloading the entire database results in other challenges for the average user, especially with the download size for the gnomAD database approaching 100GB. EXtrACtor allows the user to supply a list of genes, transcripts or regions to be queried against either the ExAC or the gnomAD database. At the same time it gives the user a flexible user interface in order to quickly filter on various properties such as cohort, variant type and more.

**Availability and implementation:** EXtrACtor source code is freely available from https://github.com/dagrim1/extractor and is written in platform independent software Java for use on windows, UNIX and mac.

## 1. INTRODUCTION

The Exome Aggregation Consortium (ExAC) is a data set based on the whole-exome sequencing (WES) data of over 60,000 unrelated individuals sequenced as part of a number of disease-specific and population genetic studies (ref). The Genome Aggregation Database (gnomAD) expands on this by increasing the number of whole-exome sequences (WES) to more then 123,000 while also including over 15,000 whole genome datasets (WGS, ref). Together, the individuals in these datasets have largely been selected for one of several common diseases and individuals affected by a severe pediatric disease have been removed. Therefore, it is assumed that these databases can be used to assess the frequency of relatively rare genetic variation in genes that are associated with rare and severe disorders in unaffected people. Such queries in these databases are used by researchers to predict pathogenicity of single putative disease causing variants observed in people affected by a severe disorder, or to test for purifying selection for Loss of Function (LoF) or missense mutations. Further, researchers may aim to test for an enrichment of rare variation in a single gene, or in a set of genes in selected patients that suffer from a rare and often severe condition. The comparison of a rare variants detected by WES or WGS in patients versus the variants listed in ExAC and gnomAD has been criticist for the inherent bias of population stratification and uncontrollable differences caused by differences in sequencing quality. However, such bias can be to some extend be controlled for by comparing the rate of variation in the test gene(s) versus several sets of control genes, which can be easily extracted using the presented tool. Within ExAC and gnomAD, the data has been harmonized and reprocessed through the same pipeline to ensure maximum consistency and uniformity across the various projects and thus removing possible batch effects and artifacts as a result of different methods and tools being used for processing and analysis of the data. It can therefore serve as reference data set for other studies, especially those that aim to detect rare disease variants.

The ExAC and gnomAD data are freely available through a browser interface and as downloadable data files. The browser can be used to search for a single gene, region, transcript or (multi-allelic) variant after which information will be presented for the canonical transcript of this gene. This information contains a coverage plot, a table containing a list of variants with their respective information concerning allele frequencies, chromosome and position, consequence, allele counts, frequencies and more. From here on the user can click on variants to look into them in more detail, or choose different transcripts for the gene of interest. However, it is only possible to search for a single entry at a time. The results can then only be filtered in a very basic manner meaning the user would have to export the results returned in order to do further and extended processing of the data. The limited search options and post processing steps makes data extraction for multiple search terms repetitive, labor intensive, and time consuming.

Multiple extractions are increasingly performed, as a researcher rarely is interested in a single gene and in general will need to scan multiple candidate genes, or regions, for a single study of one disease. The main basic aim of such studies is to filter out all putative disease variations detected for the studied rare disease that are also present in ExAC and/or gnomAD. Furthermore, ExAC and gnomAD data is also used to assess the type and frequency of aminoacid changing mutations to make statements about the probability of detected rare disease mutations. On top of that, in rare, but increasingly in common disease, multiple gene burden test are now popular approaches to overcome lack of power, especially in diseases that are characterized by a high degree of genetic heterogeneity. For such studies it is necessary to perform multiple extractions, extract additional data, and have tools for quantification and processing of extracted data. Other applications need further complicated requests to explore the ExAC and gnomAD data to a deeper extend. For example when specifically looking at certain populations and only looking at unique or unknown variants without a reference SNP cluster ID, an rs-number.

The ExAC and gnomAD developers have also made the data available for download allowing the user to directly access it. However, this introduces further complexities including handling of a number of separate files, including but not limited to VCF files of variant sites, per basepair coverage data, cnv counts and gene intolerance scores. Furthermore, the size of in particular gnomAD of almost 100GB limits the use of general programs to query data. Handling of such data require extensive bio-informatics skills or resources in order to extract the information researchers need.

EXtrACtor, which is built to answer these requests by our researches, solves the above difficulties in the form of an application allowing the user either to upload a list of either gene-names, locations or transcripts or enter them manually when querying just a single term, or limited number of terms. It then queries the ExAC or gnomAD website for the data which would normally be returned in the browser (variation data, coverage information, etc) and after the data is retrieved quick selections can be made on which exact filtering steps can be executed, updating the data in EXtrACtor in real-time without the need to generate new queries to the ExAC database. Finally, at any time the data can also be exported from the application for further use in the form of text or excel files.

## 2. IMPLEMENTATION

EXtrACtor is written in the Java language and as a result can be used across platforms. It is tested on Windows, Unix and MacOS. The graphical user interface means that no informatics background is necessary for the user to work with it, selections can simply be made using this user interface.

EXtrACtor makes use of the GNU Trove library (Trove, 2014), which supplies a more light- weight implementation of the java.util.Collections API and thereby improves on memory usage and speed, as well as the Jackson-json Parser library (Jackson, 2017) which is a library to convert JavaScript Object Notation (JSON) strings to JAVA objects and vice versa.

The applications sends a HTTP GET request to the ExAC or gnomAD database, similar to standard internet browsers, and this will return a piece of HTML code which is normally used to display the webpage in the user’s browser.

However, the actual data displayed relevant for the researchers is present in this HTML page in the form of various sections of JSON data. These sections can be found by looking for the specific keywords always preceding them, where each specific data subset such as variant information and coverage data has its specific keyword, and can thus be extracted from the HTML code in this way. The JSON data is basically a mapping of keys and their respective values.

Using the Jackson parser API the extracted JSON parts can then be easily processed after which all the entries contained in them are available to the application, meaning they can be used for calculations immediately. The full dataset is stored in the applications memory during the run, which allows the user to make changes that are processed immediately without the need to rerun a request. Therefore, variant counts can be updated right away based on the specific variant selections made by the user, but also values such as frequencies can be recalculated depending on the specific populations selected, as this information is contained in the full dataset. These features allow the user to immediately see the results of their selections making it very easy and flexible to explore the data in any way he or she desires.

At any time the user can save the currently displayed data to a tab delimited text or excel file by using the save function in the application itself, allowing the data to be used in other tools as well. It is also possible to store cached data for the various searches done, this will prevent the need to perform the same query too often and can increase speed of repetitive searches tremendously. At any time the application can be told to query the data again from the website, optionally overwriting the existing cached information.

## 3. FEATURES / CONCLUSION AND RESULTS / DISCUSSION

One of the main advantages of EXtrACtor is the possibility to query a larger number of search terms by selecting a file containing a list of these terms. The application will then automatically retrieve all the ExAC or gnomAD data for these variants and when done display them in the application’s main screen in a tab containing a table of variants. This will also show the total number of results found using the current filter settings specified by the user. Once the data has been retrieved the user can start post-processing it.

a. **Variants** The ‘Variants’ tab contains the list of variants as returned for the specified search terms, it contains all of the information as shown on the website as well as some additional ones such as population frequency data, something requiring an extra action on the side of the user on the webpage in the form of selecting the variant the user would like to see this information for. This tab also allows users to explore the data using the extensive filtering possibilities available, which are explained next.
  i. **Filtering** Various filtering selections can be made where the displayed table will be updated right away. Counts or frequencies will be recalculated using the complete dataset that is available in the memory of the program. Filters are divided in a few categories as a simple checkbox allowing the user to include or exclude that specific property. The first category is the Global category containing a dropdown box allowing the user to specify the variant type such as Loss of Function (LoF) only variant, LoF and missense variants or simply all of the variants. Subsequent filtering possibilities are the in- or exclusion of unique variants, variants with a rs-id, variants actually containing an annotated consequence, indels, or variants having their source in either genome or exome data, all these and those in the following categories are simply checkboxes. A more deep filtering can be done on the annotation level, allowing the user to select and deselect specific variants such as 3 prime UTR variants, downstream gene variants, frameshift variants, splice acceptor variants, etc. This list is generated based on the actual data returned to the application and thus is no fixed list. This ensures only relevant filters are shown to the end user. There is also the possibility to specify the various populations a researcher is interested in. The user can select any combination of populations and also specify if the variant should be present in any or all of those. Based on the selected populations frequencies and counts for the variants will be adjusted to reflect the selected populations.
  ii. **Search and quick-filter** While the default filtering exists of a number of fixed selections and options there is also a way to quickly search or filter based on user entered data. When a person wants to quickly look for variants containing a certain keyword, or in a specific region, the user can also directly enter these search terms in a row displayed above the table containing the data variants. Comparators such as greater or less then (>, <) are recognized depending on datatype displayed in the corresponding column. Queries can be made that match part of texts, anything starting or ending with a certain text, etc. This can be specified for each column separately allowing the user to do more advanced filtering such as for example: “all LoF and missense variants between base pair 145000 and 146000 having a Gly to Ala amino acid change with coverage above 50x in the exome data and coverage above 35x in the genome data.” Such a search is impossible to directly on the gnomAD website and would require the user to first download the entire variant list and then do this manually in for example Microsoft Excel.
b. **Other info** The ‘Other info’ tab shows more general info on the search term, for example mean coverage for the gene as well as a description of the gene retrieved from
c. **Feature info** The ‘Feature info’ tab will show information on the features of a gene, when a gene searches are performed. It will list the various exon, UTR and CDS elements, and their starting and ending locations, but also the largest empty space in basepairs and percentage of the feature. This can be of interest when for example a novel variant is found by the researcher in a genomic location that is void of reported variations in ExAC and gnomAD.
d. **Gene Visualization** The ‘Gene visualization’ tab displays a visual representation of the genes and variants found. In case multiple genes are queried the tab will display a list where genes can be selected for display. This functionality is limited for gene searches only, as these will also return the relevant feature data, which is not returned for region or variant searches. The display allows saving of the image to .png format, zooming in and out and it shows information on the feature or variant currently selected by the user by means of the mouse. Another possibility is to load a list of variants into the view, these will be displayed above the gene and variants retrieved from ExAC and gnomAD and allow for a comparison with an external set when desired. All the tabs are using the same base dataset, meaning all filters applied and changes made will be reflected in all these various views.

## 4. Use case

The simplest example would be a researcher having an interest in a certain disorder and has a number of variant candidates from a WGS experiments ranging across a large number of genes.

This researcher can simply select a file containing a list of these genes in the tool after which the tool will retrieve all variants for the specified genes from the gnomAD or ExAC website in order to allow the researcher to do further analysis on the variants. The specified format of the genes can be either gene names or ensemble gene ids.

Another example could be a researching searching for a list of specific variants in a gene, for example a query could be:

*All unique missense and LoF variants in the PCSK9 gene after basepair 55509607 having the PASS filter and having no rs-id while not being a 3 prime UTR or 5 prime UTR variant and being present in the European (Non-Finnish) population having a C reference allele.*

This can be quite easily retrieved using a combination of the standard and quick-filters as can be seen in the screenshot below.

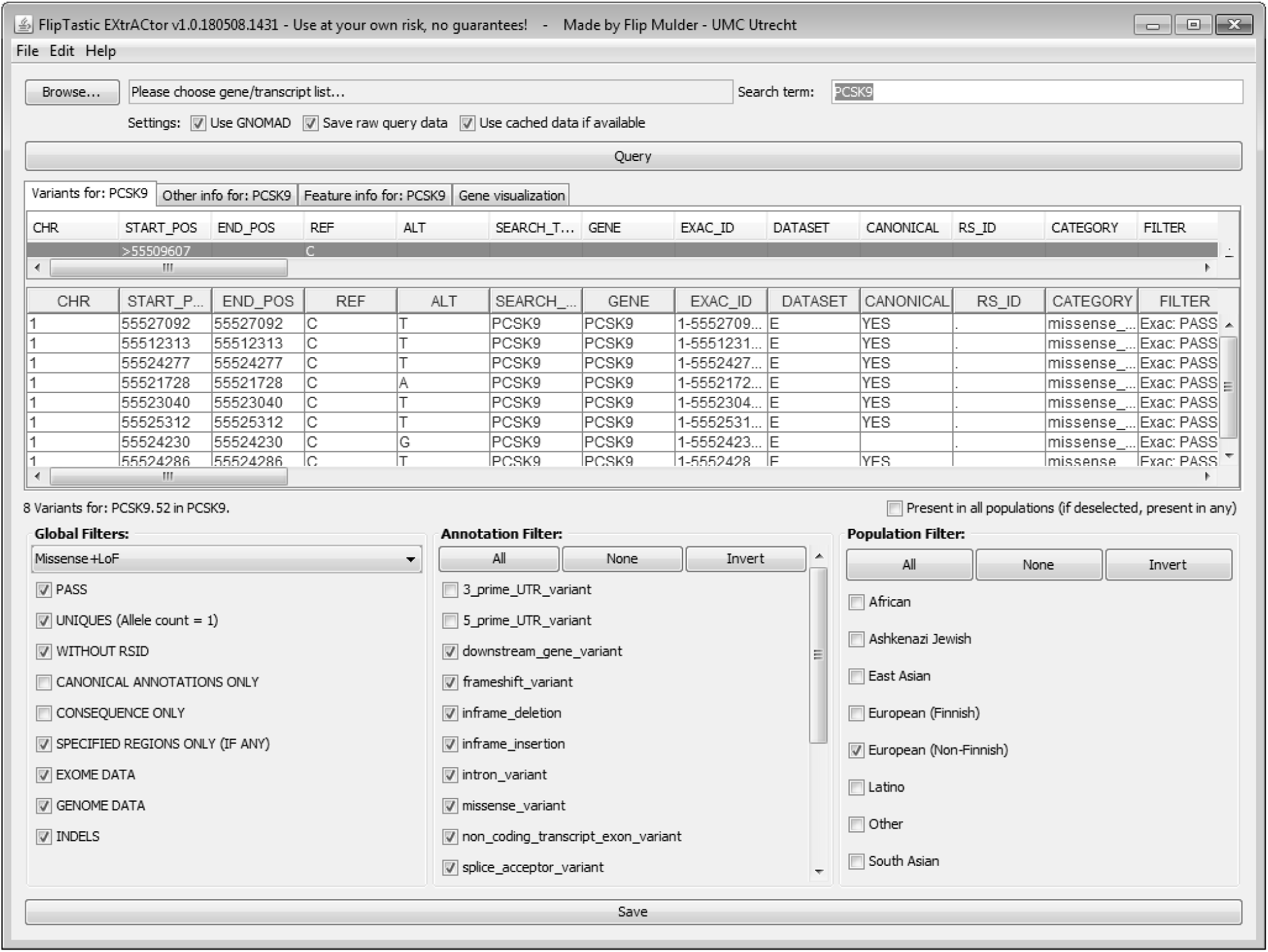

As can be seen by the line below the table, this results in a number of 8 variants for this specific search case. The 52 count shown right after it is the count not taking into account the quick-filter, so the count as resulting applying just the main filter.

To further inspect the distribution of the variants across the gene the user can switch to the Visualization tab, the following image shows this visualization tab, which gives a graphical representation of the locations of these variants in the gene. The variant types such as LoF, Missense+LoF and others are also color coded, giving a direct impression of the location of the various variants.

As can also be seen some details can be shown by hovering over a variant with the mouse, this will in turn display the ExAC id, annotation and consequence.

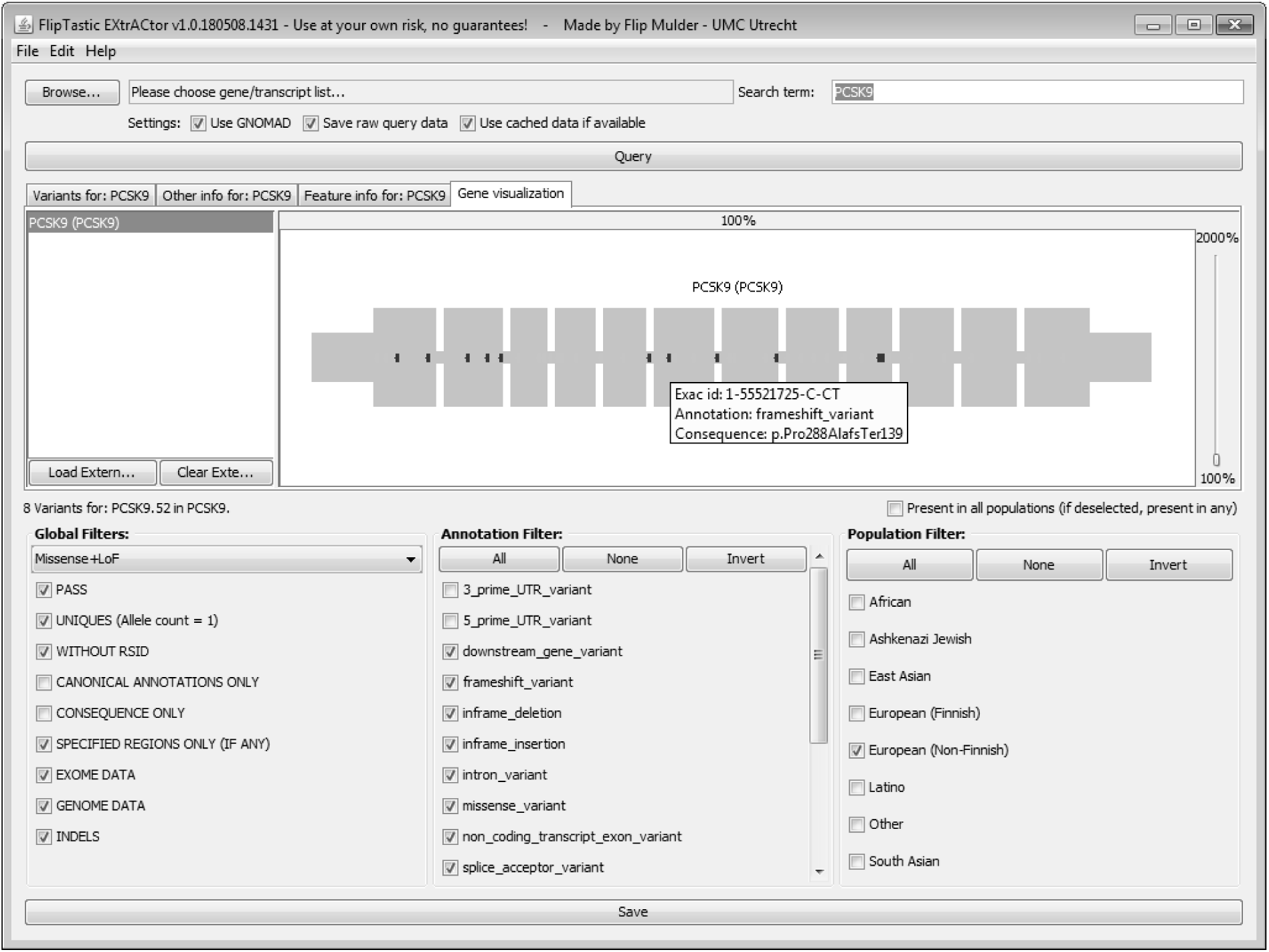

We believe this tool can greatly assist in exploring the extensive amount of data available through the ExAC and gnomAD websites.

## ACKNOWLEDGEMENTS

The authors would like to thank their colleagues at the UMC Utrecht – Genetics Department for the extensive testing, bug reports, usability input and feedback on what should and should not be possible in the program.

The authors would like to thank the Exome Aggregation Consortium and the groups that provided exome variant data for comparison. A full list of contributing groups can be found at http://ExAC.broadinstitute.org/about and http://gnomad.broadinstitute.org/about.

